# Anti-EGFR Fibronectin Bispecific Chemically Self-Assembling Nanorings (CSANs) Induce Potent T cell Mediated Anti-Tumor Response and Downregulation of EGFR Signaling and PD-1/PD-L1 Expression

**DOI:** 10.1101/2020.04.22.054338

**Authors:** Ozgun Kilic, Marcos R. Matos de Souza, Abdulaziz A. Almotlak, Jill M. Siegfried, Carston R. Wagner

## Abstract

Numerous approaches have targeted the Epidermal Growth Factor Receptor (EGFR) for the development of anti-cancer therapeutics, since it is over-expressed on a variety of cancers. Recently, αEGFR chimeric antigen receptor (CAR)-T cells have shown potential promise for the immunological control of tumors. Our laboratory has recently demonstrated that bispecific chemically self-assembled nanorings (CSANs) can modify T cell surfaces and function as prosthetic antigen receptors (PARs). This technology allows selective targeting of tumor antigens due to high avidity of the multimeric rings, while incorporating a mechanism to dissociate the rings to prevent further T cell stimulation. Previously, PARs with single-chain variable fragments (scFvs) have been successful *in vitro* and *in vivo*, activating T cells selectively at the tumor site. Alternatively, here we report fibronectin (FN3)-based PARs with improved properties such as increased protein yield, rapid protein production, increased protein stability and predicted low immunogenicity due to the human origin of fibronectins. We examined the cytotoxicity of EGFR-targeting PARs *in vitro* in which the affinities of the αEGFR fibronectins, the αEGFR/ αCD3 valency of the CSANs and the antigen expression levels were varied. Based on these selective *in vitro* cytotoxicity results, we conducted an *in vivo* study of FN3-PARs using an orthotopic breast cancer model. The FN3-PARs demonstrated potent tumor growth suppression with no adverse effects. Furthermore, these results demonstrated that FN3-PARs modulated the tumor microenvironment by downregulating EGFR signaling resulting in decreased PD-L1 expression. In addition, the expression of PD-1 was also found to be reduced. Collectively, these results demonstrate that FN3-PARs have the potential to direct selective T cell targeted tumor killing and that αEGFR FN3-PARs may enhance anti-tumor T cell efficacy by modulating the tumor microenvironment.

## INTRODUCTION

Cancer immunotherapy has gained wide spread interest over the last decade due to the promise of potent, selective and long lasting eradication of tumors. The common formats that have been developed for T cell redirecting immunotherapies are Chimeric Antigen Receptors (CARs) and bispecific antibodies. CAR-T cells are generated by genetically modifying a patient’s T cells to express a protein targeting a tumorspecific antigen (1). In contrast, bispecific antibodies generally incorporate at least two domains, one targeting the T cell and the other targeting the tumor antigen of interest resulting in T cell redirection to the tumor antigen expressing cells (2,3).

Recently, two CAR-T cell therapies have been clinically approved for B-cell lymphomas (4–8). However, for solid tumors, only modest clinical success has been observed for CARs due to a number of reasons, including CAR-T cell exhaustion, antigen loss, insufficient tumor penetration and toxicity toward normal tissues (9). Traditionally, single-chain variable fragments (scFvs) have been used as the tumor-targeting ligand for CARs. However, recently engineered protein scaffolds have been utilized to design CARs with improved properties and comparable efficacy to scFv-based CARs (10–13).

Engineered protein scaffolds have several advantages over traditional scFvs (14). Generally, they are smaller in size, have high stability and are easier to recombinantly produce and isolate (15). The protein scaffolds such as fibronectins, affibodies and DARPins are amenable to modifications and have been used in various therapeutic applications such as kinase modulation, drug delivery and immunotherapy (16–21). Fibronectins derived from the tenth type III domain of human fibronectin exhibit low immunogenicity and thus have been investigated for their immune cell targeting and tumor imaging (10). Analogous to the complementarity-determining regions of antibodies, the flexible loops connecting the seven β-strands of fibronectins can be engineered to provide target-specific binders with high affinity and specificity (22). One such target, epidermal growth factor receptor (EGFR), is a well-studied cancer biomarker and an established target for cancer therapeutics (23,24). A series of EGFR-targeting fibronectins differing in affinity and stability have been discovered by mRNA display previously (25). CAR-T cells expressing one of these EGFR-targeting fibronectins have shown *in vitro* and *in vivo* efficacy (10). When compared to cetuximab scFv CARs, FN3 and affibody CARs were able to similarly reduce the growth of xenograft tumors (10). Additionally, the FN3 and affibody CARs demonstrated enhanced efficacy *in vitro* and *in vivo* when the CARs were engineered with dual-targeting ligands (11). In another study, HER2-targeting DARPin CARs displayed improved cytotoxicity compared to scFv-based CARs (12). Similarly, VEGFR-2 targeting FN3 CARs demonstrated cytotoxicity to VEGFR-2 positive cells when they were expressed in Jurkat and primary T cells (13). Collectively, these studies suggest that alternative protein scaffolds can be deployed in CAR-T cells with at least similar efficacy to that observed for traditional scFv-based CARs.

Our group has investigated the use of prosthetic antigen receptor (PAR)-T cells. PAR-T cells are prepared by incubation of T cells with bispecific Chemically Self Assembling Nanorings (CSANs) that afford multimeric bispecific constructs incorporating an anti-CD3 scFv and a tumor targeting scFv (26). The multivalency of the CSANs enables functionalization of T cells via multiple binding interactions to CD3 on the cell surface with high stability both *in vitro* and an *in vivo* (27). The enhanced avidity of the rings provides enhanced binding and selectivity to T cells and the targeted antigen of interest (28). We have successfully demonstrated the *in vitro* and *in vivo* efficacy of scFv-based PARs (26,29). However, generating scFv-based monomers to form the rings is dependent on refolding steps that increases the time required for purification with significantly reduced protein yields (29,30). In contrast, the FN3-based monomers used for preparation of bispecific CSANs lack cysteines and thus can be expressed as soluble proteins by *E. coli* (25). We demonstrate that CSANs prepared from αCD3-scFv-DHFR^2^ and αEGFR-FN3-DHFR^2^ monomers are highly stable and enable the exploration of targeted affinity tuning of the associated PAR-T cells to EGFR (31). In addition, we investigated the anti-tumor potency of αEGFR/αCD3 PAR-T cells both *in vitro* and *in vivo*.

## RESULTS

### Preparation of FN3-1DD Fusion Proteins

Four EGFR-binding fibronectins were generated with mRNA display and their sequences have been previously reported (25). Of the four EGFR-binding fibronectins, the clones E1 (K_d_ = 0.7 nM) and E4 (K_d_ = 0.13 nM) were selected due to their lack of cysteines. Cysteine-free proteins generally result in faster purification, higher protein yields, and the potential to incorporate site-specific cysteines for further conjugation (14). E1 and E4 were fused to the N terminus of the DHFR^2^ (1DD) construct, yielding, E1-1DD and E4-1DD. Both fusion proteins were expressed in *E. coli* as soluble and monomeric proteins and Size Exclusion Chromatography (SEC) data confirmed the monomeric nature of E1-1DD (Supplementary Fig. S1A) and E4-1DD (Supplementary Fig. S1B). Similar to previous studies, upon incubation with Bis-MTX ring formation was readily observed. (Supplementary Fig. S1A and B).

For T cell targeting, we have previously designed and utilized an αCD3-scFv-DHFR^2^ fusion protein (1DD-CD3). 1DD-CD3 contains a one amino acid glycine linker between the two DHFRs and a 15-amino acid glycine-serine linker between the C-terminal DHFR and the αCD3-scFv, derived from the αCD3 monoclonal antibody, UCHT1 (27). Additionally, a low affinity EGFR peptide (K_d_ > 1 μM) (32) fused to 1DD construct, 1DD-EGF, was also expressed as a soluble protein in *E. coli* for comparison with E1-1DD and E4-1DD.

### Fibronectin CSANs showed binding to EGFR-positive cancer cells and internalization

E1-1DD and E4-1DD showed selective binding to EGFR-positive cancer cells, in correlation with the receptor densities of the cells (Supplementary Fig. 1C-E). The affinity titration experiments demonstrated high affinity towards EGFR for both E1-1DD (K_d_= 67.3 nM) and E4-1DD (K_d_= 26.3 nM) (Supplementary Fig. S1F and G). We also observed binding of E1-1DD and E4-1DD to the mouse mammary cell line HC-11 (Supplementary Fig. S2A and B). Consistent with the rapid internalization of EGFR, fluorescence microscopy studies demonstrated that E1-1DD monomers and resulting E1-CSANs undergo endocytosis by the EGFR-overexpressing epidermoid carcinoma cell line. At 37°C, both the monomers and CSANs were found to be internalized, while binding to the cell membrane was observed at 4°C. (Supplementary Fig. S3A-F).

### EGFR Downregulation of E1-1DD

EGFR signaling is oncogenic and detrimental to cancer cell survival (23). The anticancer activity of TKI and mAb therapies typically relie on the prevent this signaling pathway (33). Previously, E1 has been shown to inhibit EGFR phosphorylation with IC_50_= 38 nM against A431 cells (25). We investigated whether this effect was retained in the E1-1DD fusion construct. Compared to the control PBS group, a significant decrease in phospho-EGFR along with a decrease in the phosphorylation of downstream targets, AKT and ERK 1/2 was observed for E1-1DD treated MDA-MB-231 cells (Supplementary Fig. 4A). Nevertheless, the downregulation of receptor signaling was not accompanied by an increase in cytotoxicity. Even at high concentrations (8μM), E1-1DD treatment had no effect on MDA-MB-231 cell proliferation (Supplementary Fig. 4B).

### Bispecific CSANs bound both to cancer cells and T cells

Previously, we have demonstrated that bispecific CSANs can be prepared by mixing a 1:1:1 ratio of the chemical dimerizer, Bis-MTX and two different antigen targeting DHFR^2^ monomers (27). Similarly, bispecific αEGFR/ αCD3 CSANs were generated by mixing Bis-MTX with equal molar amounts of 1DD-CD3 and either E1-1DD or E4-1DD. The bispecific nature of the αEGFR/ αCD3 CSANs was confirmed by checking binding of the rings to two cell lines where each cell type is positive for either CD3 or EGFR (Fig. 1A). Hence, binding of the rings to both the T cells and MDA-MB-231 cells indicates that the rings contain both αCD3 and αEGFR targeting ligands. In addition, as determined by size exclusion chromatography (SEC) (Fig. 1B) and dynamic light scattering (DLS) (Fig. 1C), the hydrodynamic radius and size distribution of the bispecific rings were found to be similar to the values previously observed for CSANs (26,28,29).

**Figure 1.**
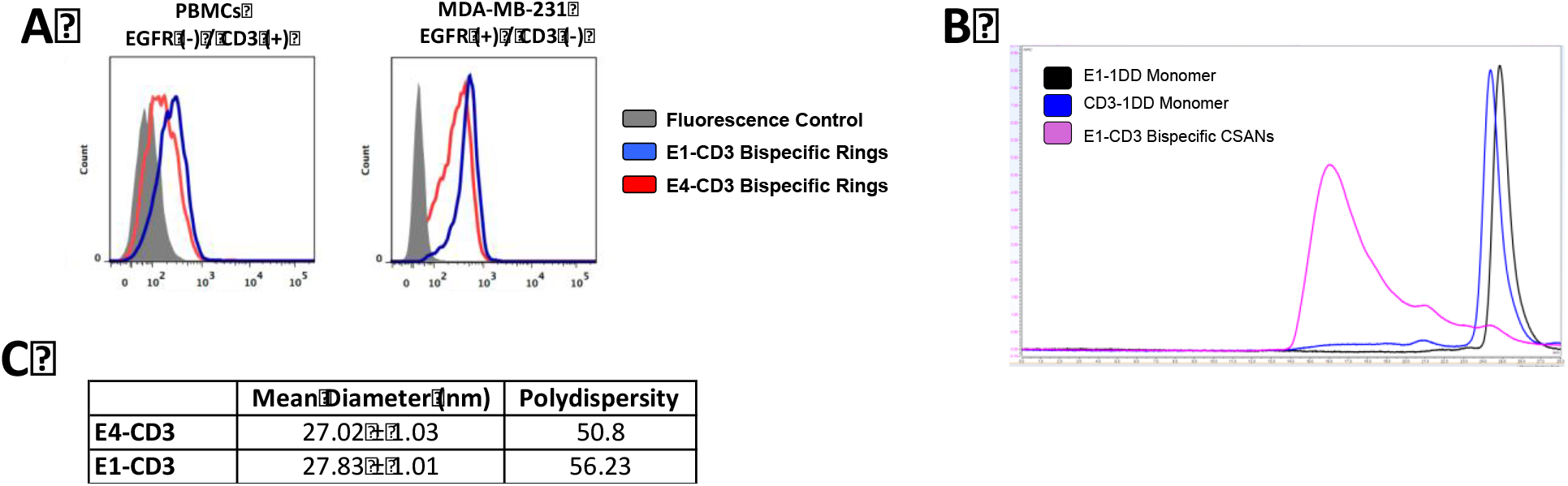
Bispecific CSAN Formation and Characterization. **A.** The bispecific nature of CSANs was confirmed by binding to two cell types. Effector T cells (left) are only CD3 positive and cancer cells (right) are only EGFR positive. **B.** The monomeric nature of E1-1DD and CD3-1DD were confirmed with Size Exclusion Chromatography (SEC). Furthermore, SEC was useful with the ring size distribution **C.** DLS results showing the mean diameter and polydispesity of the bispecific rings. **D.** Binding of both E1-1DD and E4-1DD to mouse mammary cell line (HC-11) indicating these fibronectins are mouse cross-reactive.

### Controlled disassembly of the bispecific CSANs was obtained with trimethoprim

A common concern with current immunotherapy approaches is the potential for over-activation of the T cells leading to potentially life threatening cytokine release syndrome (1). CSANs have a major advantage over these treatments because CSANs can be disassembled by addition of an FDA-approved drug, trimethoprim, thus preventing overstimulation of the T cells. Previously, we have demonstrated that trimethoprim treatment leads to disassembly of the CSANs, disruption of bispecific CSANs-induced cell-cell interactions and reduction in the levels of IL-2, IL-6, IFN-γ and TNF-α *in vivo* (26,29). Similarly, both E1 and E4 αCD3 bispecific CSANs could be dissociated with the addition of trimethoprim in a concentration dependent manner (Supplementary Fig. S5 A and B).

### αEGFR-FN3 PARs were selectively cytotoxic to EGFR-high cell lines

Besides being highly expressed on cancer cells, EGFR is also present on normal tissues, particularly the epidermis (34). Therefore, selectivity towards higher EGFR expressing cancer cells is crucial to avoid toxicity (35). We investigated the cytotoxicity of the αEGFR/ αCD3 CSANs to cell lines with variable EGFR expression levels. We determined the cytotoxicity of the E1-1DD and E4-1DD based αEGFR/ αCD3 CSANs toward the cell lines A431 (epidermal, 13 × 10^6^ EGFR per cell, 100%), MDA-MB-231 (TNBC, 0.7 × 10^6^ EGFR per cell, 5.38% relative to A431) and MCF-7 (breast, 0.2 × 10^5^ EGFR per cell, 0.15% relative to A431). These results were compared to those for a low affinity EGFR-binding peptide (pEGF-1DD, K_d_ > 1 μM) based αEGFR/ αCD3 CSANs developed previously by our laboratory (32). No significant toxicity was observed for each of the αEGFR/ αCD3 CSANs toward the lowest EGFR expressing cell line, MCF-7 (Fig. 2A). In contrast, significant and comparable cytotoxicity was observed for each of the E1-1DD and E4-1DD based αEGFR/ αCD3 CSANs toward higher EGFR expressing MDA-MB-231 (Fig. 2B) and A431 cells (Fig. 2C). In both cases, maximum cytotoxicity was observed after treatment with 12.5 nM of the CSANs. Similarly, the cytokine profiles observed for the treatment groups correlated with the levels of cytotoxicity, with significant production of IL-2 and IFN-γ observed for the treatment of MDA-MB-231 cells with E1-1DD and E4-1DD based αEGFR/ αCD3 CSANs, compared to MCF-7 cells (Fig. 3A-D). As observed for induced cytotoxicity, the maximum cytokine production occurred after treatment with 12.5 nM of the CSANs. These results suggest that αEGFR/ αCD3 CSANs induced cytotoxicity is dependent on the affinity of the αEGFR ligand and the cell surface expression levels of EGFR.

**Figure 2.**
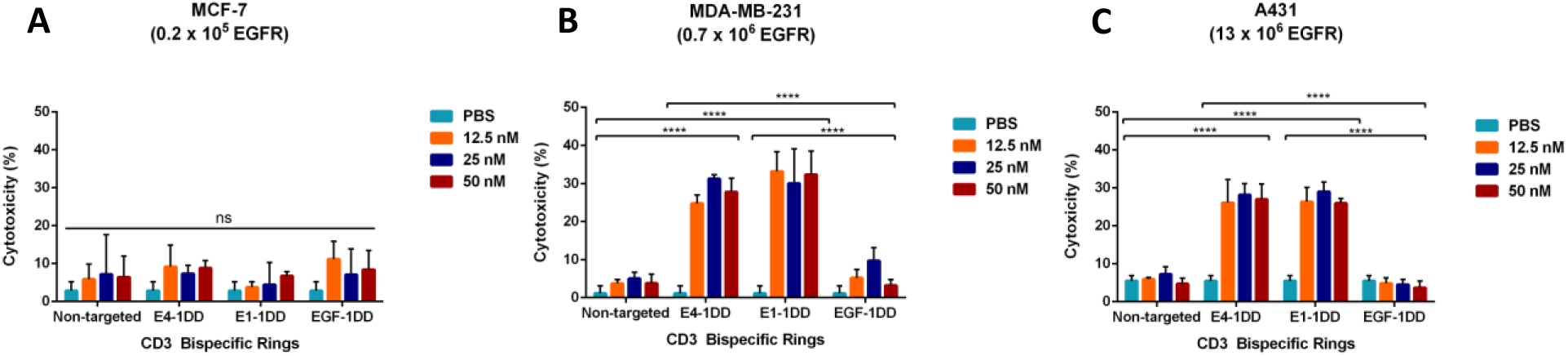
Cytotoxicity of EGFR-targeting PAR-T cells. **A.** Cytotoxicity toward MCF-7 cells **B.** Cytotoxicity toward MDA-MB-231 cells **C.** Cytotoxicity toward A431 cells. In a 96-well plate 0.5 x 10^4^ cells were seeded and incubated with the protein treatments for 24 hours. The supernatants were analyzed for cytotoxicity using LDH assay. *P<0.05 with respect to non-treated controls, by one-way ANOVA analysis.

**Figure 3.**
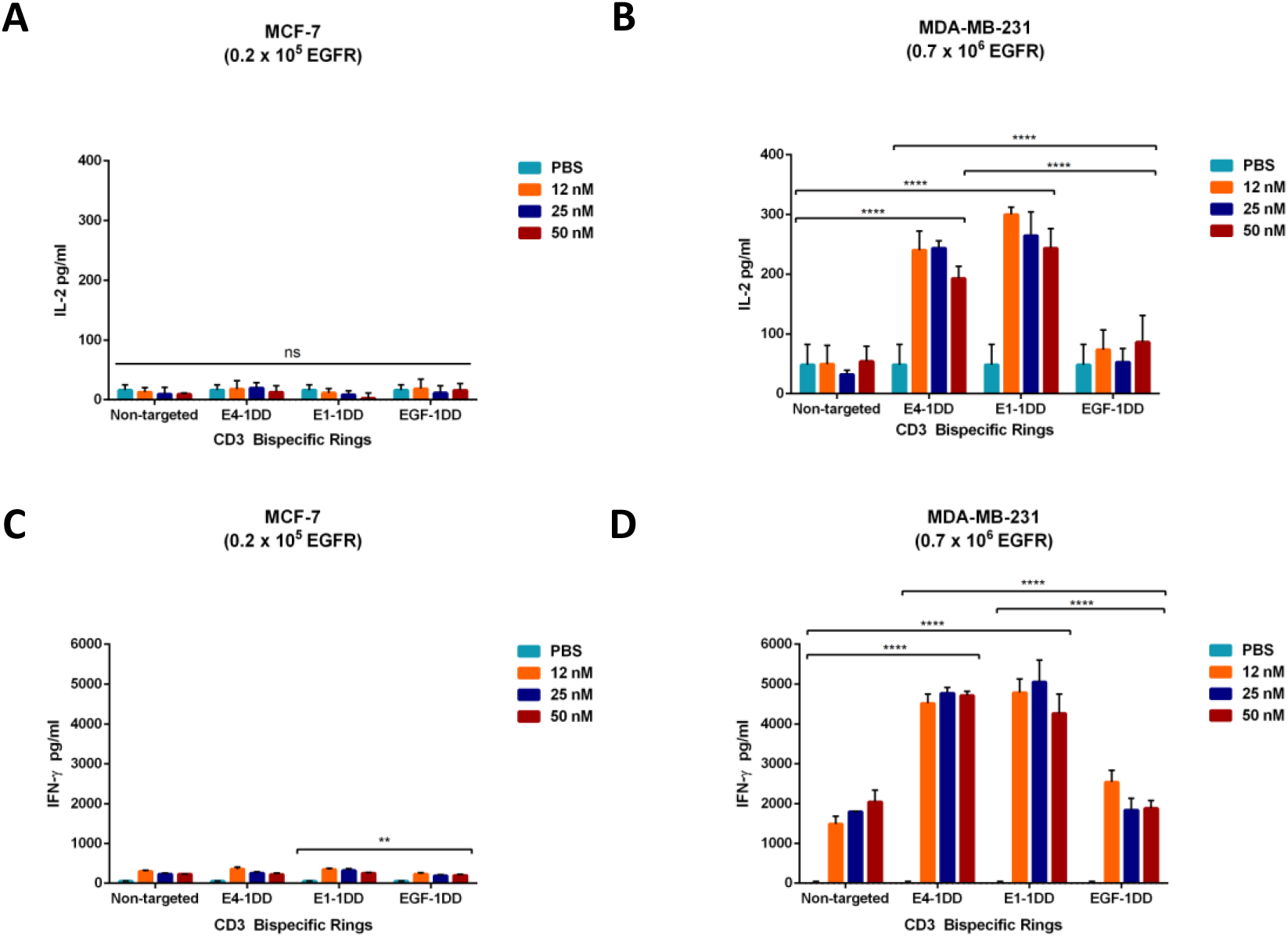
Cytokine Release Profile of EGFR-targeting PAR-T cells. **A.** IL-2 released from MCF-7 cells after cytotoxicity study **B.** IL-2 released from MDA-MB-231 cells after cytotoxicity study. **C.** IFN-γ released from MCF-7 cells after cytotoxicity study **D.** IFN-γ released from MDA-MB-231 cells after cytotoxicity study. In a 96-well plate 0.5 x 10^4^ cells were seeded and incubated with the protein treatments for 24 hours. The supernatants were analyzed for cytotoxicity using ELISA kit. *P<0.05 with respect to non-treated controls, by one-way ANOVA analysis.

### Valency modulation of the bispecific CSANs did not enhance cytotoxicity in vitro

Previously, we reported that the distribution of each monomer in a bispecific ring could be modified by mixing different equivalences of each monomer during ring formation (29). Based on this quantitative understanding of the effects of valency, affinity and antigen expression on CSANs cell surface binding, we prepared CSANs with different monomer distributions ranging from predominantly displaying the αCD3 scFv to predominantly displaying the αEGFR FN3 (Fig. 4A). We hypothesized that a higher αCD3 valency of the rings would enhance cytotoxicity by enabling initial and preferred binding to the T cells over the tumor cells. Consequently, we prepared variable valency CSANs, by mixing variable ratios of the αCD3-1DD monomer with E1-1DD (1:7 to 7:1) and compared their cytotoxicity to other CSANs prepared with αCD3-1DD monomer and the non-ligand binding monomer, 1DD. As can be seen in Fig. 3B, varying the αCD3 to αEGFR valency with the αCD3-1DD and E1-1DD based bispecific CSANs did not affect the cytotoxicity of the bispecific CSANs. As observed before (vide supra), T cell directed cytotoxicity was observed toward the high EGFR expressing MDA-MB-231 cell line, but not the lower EGFR expressing MCF-7 cell line. (Fig. 4B and C) Thus, regardless of whether the CSANs were predominantly directed toward the T cells or the tumor cells, no significant selectivity for T cell induced cytotoxicity was observed under 2D culture conditions.

**Figure 4.**
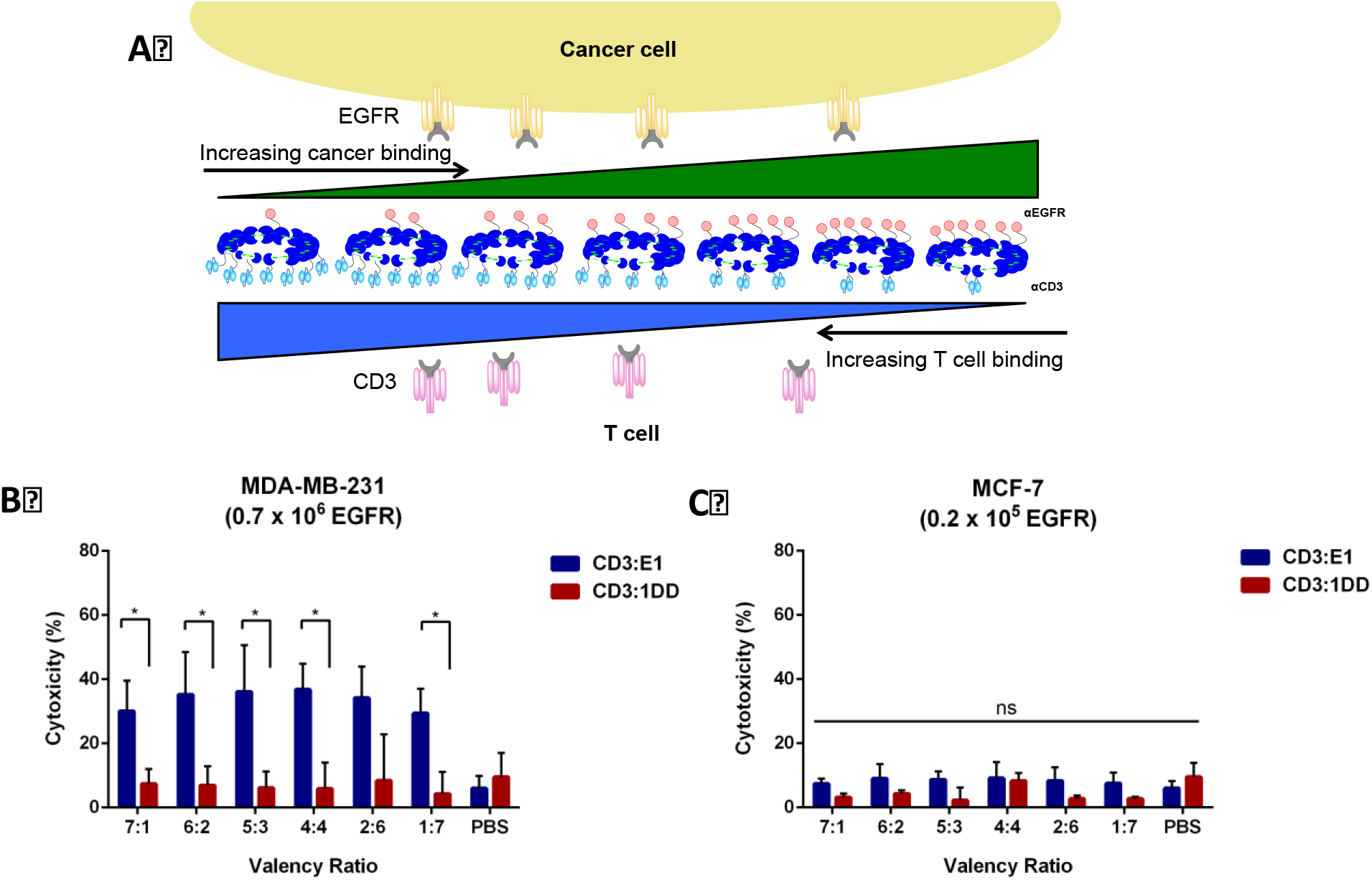
Valency modulation of bispecific CSANs. **A.** Schematic demonstrating that by varying valency of the rings, we can shift binding of rings predominantly to either cancer cells or T cells. **B.** LDH-based cytotoxicity results for MDA-MB-231 cells when valency was modulated from CD3-targeting high (left) to EGFR-targeting high (right). Other than the PBS control and 2:6 valency group, at every valency examined, E1-CD3 rings were significantly more cytotoxic compared to 1DD-CD3 rings. **C.** LDH-based cytotoxicity results for MCF-7 cells when valency was modulated from CD3-targeting high (left) to EGFR-targeting high (right). There was no significant difference between E1-CD3 and 1DD-CD3 rings at any valency ratio. *P<0.05 with respect to E1-CD3 rings and 1DD-CD3 ring controls, by 2-tailed Student’s t test.

### Bispecific CSANs showed in vivo tumor reduction and increased survival

To evaluate the *in vivo* efficacy of targeting EGFR expressing tumors with αEGFR/ αCD3 CSANs, we elected to carry out anti-tumor studies with E1-1DD based bispecific CSANs with αCD3-1DD and E1-1DD monomer ratio of 4:4 and a triple negative breast cancer (TNBC) orthotopic xenograft model. MDA-MB-231 cells were injected into the fourth mammary fat pad of young adult female immunodeficient NSG mice. Five days after the tumor induction, PBMCs were administered to the mice via intravenous (IV) injection. Once the tumor sizes reached 100 mm^3^, the mice were randomly distributed into three groups (n=5), and treated with the αEGFR/ αCD3 CSANs. The control groups were treated with either αCD3 CSANs or PBS. Each mouse received six consecutive doses (IV) every two days and the tumor size and weight of each mouse were determined throughout the study (Fig. 5A). Significant anti-tumor activity was observed after the second dose of the αEGFR/ αCD3 CSANs and continued throughout the course of the study compared to the control groups (Fig. 5B). Although the E1 FN3 cross reacts with both murine and human EGFR, no significant weight loss was observed for the αEGFR/ αCD3 CSANs treatment group (Fig. 5C). One major side effect of EGFR-based treatments is the potential for the development of skin rash due to the expression of EGFR on epidermal cells (36,37). Nevertheless, no significant development of skin rash was observed for the αEGFR/ αCD3 CSANs treated mice. Interestingly, the mice in the control groups showed symptoms of Graft-versus-Host disease (GvHD), such as weight loss and loss of fur, while the αEGFR/ αCD3 CSANs treated group exhibited no such symptoms (38). Taken together, these results indicate that αEGFR PAR-T cells have the potential to exhibit potent and non-toxic anti-tumor activity toward high EGFR expressing tumors.

**Figure 5.**
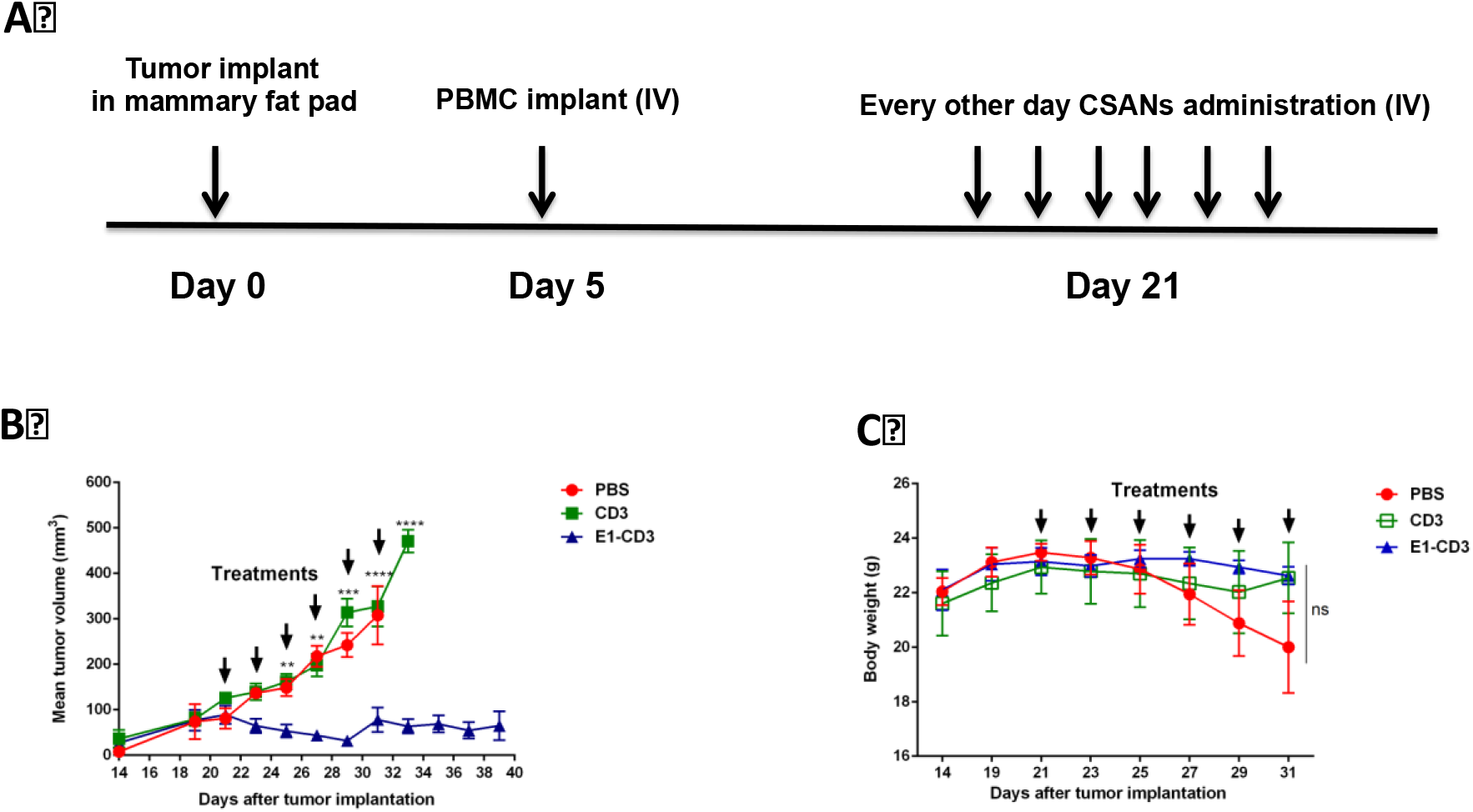
*In vivo* efficacy study of adnectin PARs using an orthotopic NSG mouse model. **A.** Schematic demonstrating the time of tumor and PBMC implantation and the treatment schedule. **B.** Mean tumor volumes were measured every other day and recorded. Tumor volume measurements were done with a caliper. Arrows indicate the days of treatment. *P<0.05 with respect to non-treated controls, by two-way ANOVA analysis followed by Dunnett’s test for multiple comparisons. **C.** Average body weight within each group is recorded. Arrows indicate the days of treatment. There was no significant difference between average weights within each group throughout the study.

### Analysis of Tumors by IHC

In order to evaluate the long-term effects of bispecific CSANs therapy, all tumors were collected and processed for Immunohistochemistry (IHC) ten days after the last treatment. H&E staining demonstrated the differential tumor cellular morphology between the two control groups (PBS, CD3) and the αEGFR/ αCD3 CSANs treatment groups (Supplementary Fig. S6A-C). While the tumors were densely packed in the control groups, the treatment group exhibited higher levels of stroma. In addition, we observed the presence of cells positive for the proliferative marker Ki67 in the treatment group, consistent with the presence of proliferating T cells (Fig. 6A).

**Figure 6.**
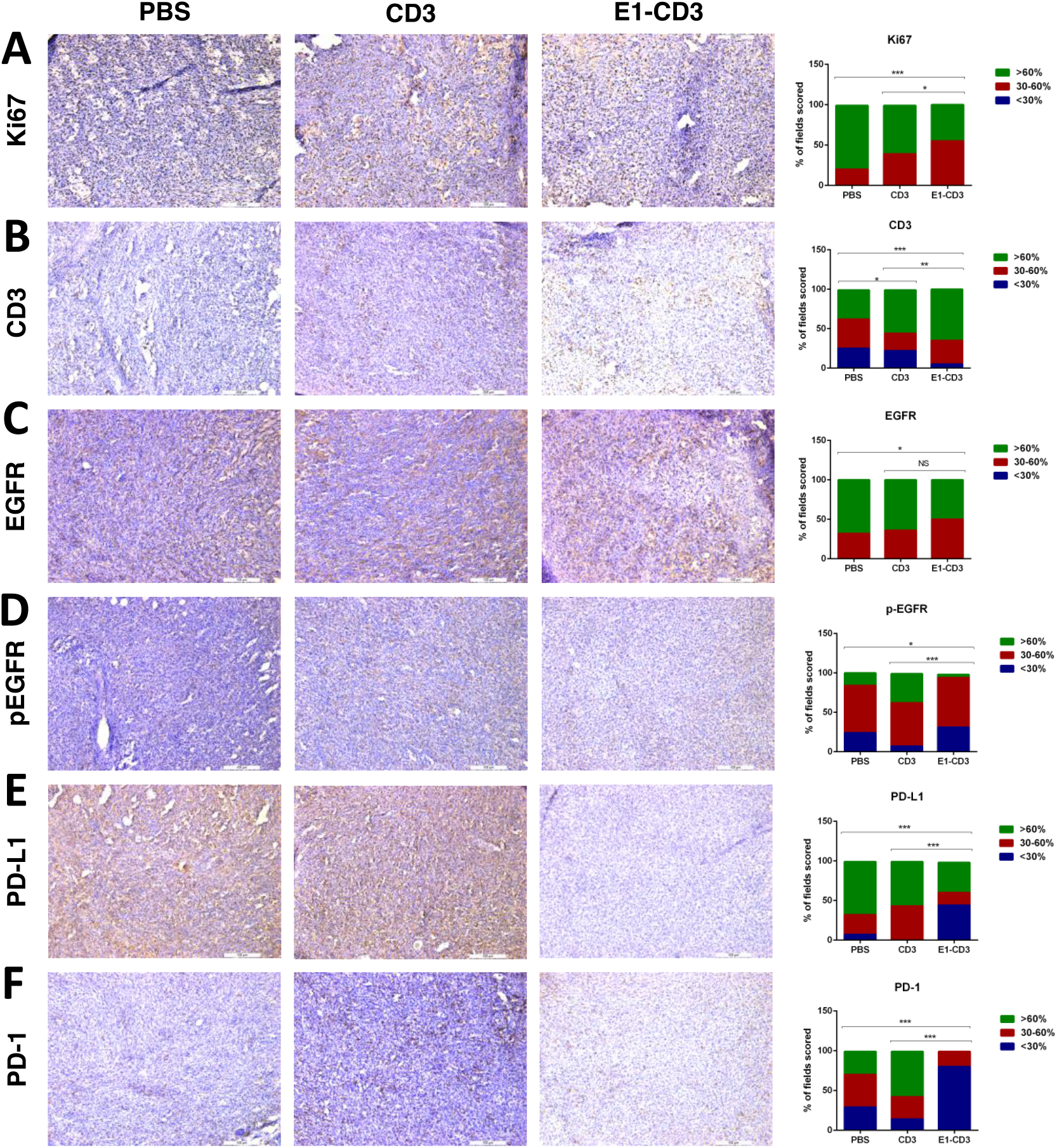
Representative Immunohistochemistry (IHC) images and quantification. **A.** ki67 **B.** CD3 **C.** Total EGFR **D.** phospho-EGFR **E.** PD-L1 **F.** PD-1. Brown staining indicates the cell is positive for the markers listed. For each staining representative images for PBS, CD3 and E1-CD3 groups are shown. On the right side of the image panels, expression of each marker is quantified where (blue) low expression, (red) medium expression (green) high expression for the marker within regions of each image.

A major concern in cancer immunotherapy for solid tumors is the inability of T cells to infiltrate into the tumor (9). T cell infiltration was observed for both the αEGFR/ αCD3 CSANs treated groups, as well as the control groups, shown by staining with an αCD3 antibody (Fig. 6B). Consequently, the injected T cells successfully proliferated in mice and survived even 10 days after the last treatment. A major hurdle in cancer immunotherapy is the loss of the targeted antigen reducing the effectiveness of immune cell targeted therapies (3). IHC analysis revealed that no significant difference in the levels of EGFR expression between the αEGFR/ αCD3 CSANs and αCD3 CSANs treatment groups and only a minor difference between the treatment groups and the PBS control group (Fig. 6C).

Since MDA-MB-231 cells express high levels of PD-L1, an immunosuppressive marker, and EGFR signaling enhances PD-L1 expression, we investigated the effect of treatment αEGFR/ αCD3 CSANs on PD-L1 (39–41). When the MDA-MB-231 tumors were examined by IHC, those that had been treated with αEGFR/ αCD3 CSANs exhibited decreased levels of phosphorylated EGFR (Fig. 6D), as well as decreased levels of PD-L1 and PD-1 expression compared to the control groups (Fig. 6E-F). The effect on this pathway was further confirmed in a parallel study in which the tumors were harvested nearly a month after the last treatment and subjected to western blot analysis (Supplementary Fig. S7A and B). While the phosphorylated and total EGFR levels were similar between PBS and E1-CD3 treated groups, the PD-L1 downregulation was significant, even 27 days after the last treatment, indicating that the bispecific CSANs had a long-term effect on immune modulation. Thus, the αEGFR/ αCD3 CSANs contributed to enhanced T cell targeted cytotoxicity, as well as a reduction in the immunosuppressive capabilities of the tumor microenvironment.

## DISCUSSION

We have previously demonstrated that bispecific CSANs can function as tumor targeting T cell engagers. αCD3/αEpCAM CSANs were shown to efficiently eradicate tumors from NSG mice orthotopically implanted with the EpCAM overexpressing breast cancer cell line, MCF-7, with no observable cytokine based toxicities (29). In addition, while immunotoxins targeting EpCAM have been shown to undergo rapid internalization, no significant internalization of αEpCAM CSANs has been observed (29). In contrast, we have demonstrated that αEGFR CSANs can undergo rapid internalization. Consequently, we decided to characterize the efficacy of T cell targeted tumor cell killing with αCD3/αEGFR CSANs, to assess the potential effect of potential rapid CSANs internalization on anti-tumor activity. Additionally, since αEGFR CSANs are cross-reactive with mouse EGFR, animal studies with this scaffold provide insights on the potential for human toxicities. Although our previous results with αCD3/αEpCAM CSANs proved efficacious, since the αEpCAM scFv binds to human EpCAM, but not the murine EpCAM, the potential for off target toxicities could not be assessed.

Triple negative breast cancer (TNBC) is an aggressive form of breast cancer with limited treatment options (33). Unlike other forms of breast cancer, which take advantage of EGFR, HER2 and HER3 signaling, TNBC cells are HER2 and HER3 negative and thus exclusively dependent on EGFR overexpression (33). Nevertheless, while tyrosine kinase inhibitors (TKIs) and monoclonal antibodies (mAbs) targeting EGFR are clinically used for lung cancer and colorectal cancer respectively, they have not shown significant clinical success as breast cancer monotherapies (33). Recently, αEGFR CAR-T cells have shown promise against solid tumors (30). CAR-T cells incorporating an αEGFR scFv have shown efficacy against gliomas, while αEGFR FN3 have demonstrated potency against lung cancer (10). Consequently, we sought to characterize the anti-tumor ability of αEGFR/αCD3 CSANs to induce T cell directed cell killing against EGFR expressing cancer cells, including TNBC.

To avoid the need for refolding and to enhance stability, we chose to prepare αEGFR/αCD3 CSANs that incorporated human FN3s, E1 (K_d_ = 0.7 nM, T_m_ = 72°C) and E4 (K_d_ = 0.13 nM, T_m_ = 69.2°C) that were shown to selectively bind EGFR (25). In addition, E1 and E4 were shown to competitively inhibit EGF binding, inhibit EGFR and ERK phosphorylation and cross react with both human and murine EGFR (25). Similar to the FN3s, the His-tagged FN3 DHFR fusion proteins, E1-1DD and E4-1DD were solubly expressed by *E.coli*, yielding 25-35 mg/L after purification by nickel affinity chromatography. In contrast, scFv based DHFR fusion monomers require an extensive refolding step, due to the inherent presence of disulfide bonds, with protein yields on the order of 1-4 mg/L.

Both the E1 and E4 monomers and corresponding CSANs were shown to bind to cellular EGFR. Similar to past binding studies with E1 and E4, E4-1DD and E4-CSANs exhibited a 2-fold greater affinity for EGFR than E1-1DD and E1-CSANs. Nevertheless, for both E1 and E4, the fusion to DHFR^2^ resulted in a nearly 85- and 200-fold reduction in binding affinity compared to FN3 E1 and E4 (25). When compared to FN3 E1 and E4, the E1 and E4 DHFR^2^ monomers and corresponding CSANs similarly underwent internalization by endocytosis (25). The binding of the αEGFR CSANs to EGFR on MDA-MB-231 cells was also found to inhibit EGFR phosphorylation. Collectively, these results demonstrated that the fusion of E1 and E4 to the DHFR^2^ monomers did not weaken their EGFR binding or impact on EGFR associated induced signaling.

For any CAR or bispecific construct, efficient immune synapse formation is critical for T cell activation, which is affected by several factors (42). T cell activation has been shown to be dependent on antigen density, with cell lines expressing high levels of EGFR, such as MDA-MB-231 (0.7 x 10^6^ antigens/cell) and A431 (13 x 10^6^ antigens/cell) were able to initiate T cell activation and cell killing when treated with E1-CSANs and E4-CSANs, while MCF-7 cells (2.0 x 10^4^ antigens/cell) were unable to initiate T cell activation or cytotoxicity with either of the respective CSANs. CAR-T cells have been shown to exhibit cytotoxicity at antigen levels as low as 200 antigens/cell while bispecific αEGFR/αCD3-CSANs require higher antigen density to initiate cell killing (43). The higher minimum antigen density required for T cell activation with either E1-CSANs or E4-CSANs, suggests that T cell activation and cell killing may have greater selectivity for EGFR-overexpressing cells, such as cancer cells, over expression on normal tissues Indeed the MCF-7 cells (2.0 x 10^4^ antigens/cell), which express similar levels of EGFR found on normal tissues (3.0 x 10^4^ antigens/cell), did not undergo αEGFR/αCD3-CSANs T-cell induced cell killing (43).

Recently, we have demonstrated both *in vitro* and *in vivo* that the binding specificity of EpCAM targeted CSANs is dependent on ligand binding affinity, valency and the antigen expression levels on the target cells (28). Consequently, we analyzed the effect of these factors on the T cell induced cytotoxicity by FN3-based EGFR-targeting bispecific CSANs. We compared the efficacy of four protein scaffolds with varying αCD3 scFv and αEGFR FN3 for EGFR. Since there was no difference in the ability of E1 and E4 to induce T cell activation and targeted cell killing, we chose to evaluate E1, since the K_d_ for E1-1DD (K_d_= 67.3 nM) and K_d_ for αCD3-1DD (K_d_= 51 nM) were the most similar and therefore would not inherently have a preference for binding to T cells or the target tumor cells (44). Despite valency variations from (αCD3)1:7(E1) to (αCD3)7:1(E1), no significant difference was observed in the efficiency of T cell induced cell killing. Thus, neither the preference of the αCD3/αEGFR CSANs for binding to the T cells or tumor cells or the potential for internalization by the tumor cells significantly altered the degree of induced cytotoxicity *in vitro*. Consequently, we chose to carry out *in vivo* studies with αCD3/αEGFR CSANs composed of an equal ratio of αEGFR FN3 and αCD3 scFv binding ligands.

Based on these results, we investigated the *in vivo* efficacy of αCD3/αEGFR CSANs on T cell induced tumor eradication with an orthotopic triple negative breast cancer xenograft mouse model implanted with MDA-MB-231 cells and human PBMCs. The bispecific rings effectively prevented tumor progression with no detectable toxicity associated with targeting EGFR, such as skin rashes and loss of body weight (36,37). Previously, CAR-T cells have been prepared with αEGFR FN3 (10,11). In one case αEGFR CAR-T cells prepared with the low affinity FN3, E3 (K_d_ = 9.9 nM) exhibited only modest suppression of the growth of NSCLC tumors with a mouse xenograft model (10). Attempts to prepare αEGFR CAR-T cells from E1 and E4 were not successful. In another study, αEGFR CAR-T cells were prepared with E1 and exhibited significant anticancer activity in a mouse pancreatic cancer xenograft model (11). Therefore, the observed anti-tumor activity of αCD3/αEGFR CSANs appears to be at least as potent as similar FN3 based CAR-T cells.

Examination of tumors for untreated and αCD3/αEGFR CSANs treated tumors revealed significant infiltration and proliferation of T cells for treated tumors, as well as tumor cell apoptosis and necrosis, compared to control tumors. Surprisingly, while the levels of EGFR expression remained unchanged, substantial suppression of EGFR phosphorylation and PD-L1 and PD-1 expression were observed several days after the last dose of the αCD3/αEGFR CSANs, compared to control tumors. While current EGFR therapies have shown efficacy against certain TNBC cells, they are ineffective against the MDA-MB-231 cells used in this study, due to the presence of the G13D KRAS mutation (33). Consistent with these observations, suppression of EGFR signaling by TKIs and targeted monoclonal antibodies has been shown to suppress tumor expression of PD-L1 (45). These observations were not described for alternative protein scaffolds, αEGFR CAR-T cells or αEGFR bispecific T cell engagers (45). Thus, treatment with αEGFR/αCD3-CSANs resulted in not only T cell proliferation and directed cell killing, but a reduction in the immunosuppressive tumor microenvironment.

Taken together, these results demonstrate the potential for employing FN3 targeted T cell engagers for cancer immunotherapy and αCD3/αEGFR CSANs in particular. αCD3/αEGFR CSANs prepared with the FN3, E1, were shown to be non-toxic and to elicit a strong EGFR targeted T cell based anti-tumor response, as well as decreasing the immunosuppressive nature of the tumor microenvironment. The low toxicity of the αCD3/αEGFR CSANs is likely due to not just the nature of the αEGFR ligand, but also on the selectivity of the bispecific nanorings for tumor cells expressing relative high levels of EGFR. As has been demonstrated before, the ability to disassemble the CSANs by dosing with the FDA approved drug, trimethoprim, offers yet one more temporally implementable safety feature unique to CSANs (46). The generality of these findings for other EGFR expressing tumors is currently under investigation.

## MATERIALS AND METHODS

### Cell Lines, Culture Conditions and T cell isolation

MCF-7, MDA-MB-231 and A431 cancer cells were obtained from American Type Culture Collection (ATCC, Rockville, MD). Cells were cultured in Dulbecco’s Modified Eagle Medium (DMEM) supplemented with 10% FBS, 100 U/mL penicillin and 100 μg/mL streptomycin at 37 °C in 5% CO2. Human PBMCs from healthy donor blood samples were isolated from buffy coats (Obtained from Memorial Blood Centers, St. Paul, MN) by Ficoll density gradient centrifugation.

### EGFR Antigen Expression Quantification

EGFR levels of the cancer cells used in this study were quantified using Quantum Simply Cellular Anti-Mouse IgG Fluorescence Calibration Beads from Bang’s Laboratories, Inc (Fishers, IN) following manufacturer’s protocol. Briefly, cells and beads were incubated with Alexa Fluor 647 labeled anti-EGFR antibody (Thermo Fisher) for one hour and after wash steps cells and beads were analyzed using BD LSR II flow cytometer and FACS Diva software.

### DHFR^2^ Protein Expression and Purification

The fusion DHFR^2^ constructs were transformed to Rosetta cells from EMD Millipore (Billerica, MA). Colonies picked were cultured in 4L of LB media, shaken at 250 rpm at 37 °C until the OD600 value reached in the ranges of 0.5-0.8 and subsequently induced with isopropyl β-D-1-thiogalacto-pyranoside (IPTG; Sigma-Aldrich, St. Louis, MO) at a final concentration of 1 mM. After IPTG induction, cultures were incubated at 37 °C for an additional three hours and centrifuged at 7500 rpm to obtain the cell pellet. 1DD-CD3 was purified from inclusion bodies as described before (27). E1-1DD and E4-1DD were purified by lysing the cell pellet in lysis buffer containing protease inhibitor and further purifying the cell lysate using a Nickel column yielding 32 mg/L and 27 mg/L, respectively.

### CSAN Oligomerization and Characterization

CSANs were formed by mixing the 1DD monomer with 2 equivalence of the C9 dimerizer molecule and incubated at 37°C for 30 minutes. The CSANs were further characterized by HPLC analysis using Sephadex column for Size Exclusion Chromatography (SEC). The hydrodynamic radius and polydispersity of the rings was determined with Dynamic Light Scattering (DLS) using pUNk by Unchained Labs (Pleasanton, CA).

### Binding Studies

Cancer cells were trypsinized and 1.0 x 10^5^ cells were incubated with 100-500 nM CSANs for one hour on ice. The cells after incubation were washed with FACS buffer and incubated with anti-his Alexa Fluor 647 antibody (Thermo Fisher Scientific) for 30 minutes on ice. Cells were again washed with FACS buffer and further analyzed with LSR II Flow Cytometer. Epitope confirmation was done by a competitive binding assay where saturating concentration of ICR-10 PE antibody (Thermo Fisher Scientific) was coincubated with a concentration gradient of unlabeled E1-1DD or E4-1DD. The decrease of fluorescence signal with increasing unlabeled protein competition was detected with LSR II Flow Cytometer.

### Internalization Studies

A431 cells were counted and plated on 8-well μ-slides from ibidi at a density of 0.2 x 10^6^ cells per well and cultured overnight at 37 °C in 5% CO2. Cells were washed and kept on ice for 2 hours followed by 1 hour incubation with the Alexa Fluor 647 stained E1-1DD monomers or CSANs. Following the treatment cells were fixed with 4% paraformaldehyde and stained with Prolong Gold Antifade mounting media. The images were deconvolved with Elements software and further analyzed with FIJI software.

### Western Blot Analysis

MDA-MB-231 cells were plated at 0.4 × 10^6^ in 6-wells plates containing DMEM medium with 10% FBS, 1%GlutaMax and 1% penicillin/streptomycin. After 24hr, the cells were washed with PBS and incubated with PBS or E1 (0.5μM) for 24, 48, and 72hr in serum free media. Whole cell lysates were collected using RIPA buffer with protease inhibitor (1X PBS, 1% nonyl phenoxypolyethoxylethanol NP40, 0.5% sodium deoxycholate, 0.1% sodium dodecyl sulfate containing 1 tablet protease inhibitor cocktail/10 mL buffer [Roche Diagnostics, Indianapolis, Indiana]). Protein concentration was determined using DC assay reagent (Bio-Rad Laboratories). Cell lysates (20 μg) from each sample was electrophoresed on 7.5% sodium dodecyl sulfate-polyacrylamide gels for 80 minutes and transferred on to a polyvinylidene difluoride membranes for 60 minutes at 100V. Membranes then were blocked with 5% milk in 0.1% TBST buffer or 60 minutes. The membranes then incubated with primary antibodies (all from Cell Signaling technology) at 1:1000 dilution. Secondary antibody was added after 3 times washing with TBST for 60 minutes. Blots then were developed using super signal West Pico according to the manufacturer’s protocol. ImageJ software was used to quantify the results.

### Cell Proliferation Assay

MDA-MB-231 cells were plated in 96-well plate at 6,000 cells/well and cultured overnight. On the second day, media were removed and E1-1DD or 1DD were added to the wells using 1% FBS basal DMEM media and PBS was used as a control. After 72 hours, cell viability was assessed using the MTS reagent CellTiter96 one solution (Promega, Madison, WI). MTS reagent (20 μL) was added to each well and incubated for 30 minutes. The microplate reader Synergy HT (BioTek) was used to read out the plate at 490 nm and Gene5 software was used to analyze the data.

### Cytotoxicity Assay

For the cytotoxicity assay 1.0 x 10^5^ cells were plated per well in a 96-well plate and cultured overnight. Next day, unactivated PBMCs were functionalized with bispecific CSANs for 30 minutes and then added to the wells containing target cells and incubated for 24 hours. The cell lysis in each well was quantified by measuring the lactate dehydrogenase (LDH) released into the media using a nonradioactive cytotoxicity assay (CytoTox 96 Non-Radioactive Cytotoxicity Assay, Promega) following the manufacturer’s manual.

### Cytokine Analysis

The supernatants were collected after cytotoxicity assay for cytokine analysis by using IFN-γ and IL-2 ELISA kits (Invitrogen). Cytokine levels were analyzed by comparing the values obtained from samples to a standard curve generated from known concentrations of the cytokines.

### *In vivo* Efficacy Study

All animal protocols were approved by the University of Minnesota Institutional Animal Care and Use Committee (IACUC) complying with both federal and institutional regulations for humane treatment of animals. On Day 0, six to eight week-old female NOD.Cg-Prkdcscid Il2rgtm1Wjl/SzJ (NSG) mice from Jackson Laboratory were injected in the fourth mammary fat pad with 1.0 × 10^6^ MDA-MB-231 cells in a 1:1 Matrigel/PBS mixture. On Day 5, 20 x 10^6^ PBMCs were injected intravenously. Mice were monitored until the tumors reached 100 mm^3^ in size and they were randomized into three treatment groups (n=5) of PBS, CD3 and E1-CD3. After randomization mice in each group received a total of 6 treatments of 1 mg/kg intravenously given every other day. Additionally, the tumor size and body weight were monitored every other day or daily if necessary. The tumor size was measured by a caliper and the tumor volume was calculated based on (length x width^2^)/2. The mice were monitored daily and endpoint was determined as significant weight loss (%15) or severe dehydration.

### Western Blot Analysis of Tumor Lysates

In a parallel study done for western blot analysis, 10 month old male NSG mice were injected in the flank with 1.0 × 10^6^ MDA-MB-231 cells in a 1:1 Matrigel/PBS mixture followed by the same treatment protocol above. Tumors were harvested 27 days after the last treatment. Tumors were homogenized and lysed using RIPA buffer and the total protein concentrations were determined using DC assay reagent (Bio-Rad Laboratories). Cell lysates (50 μg) from each sample was electrophoresed and blotted following the western blot protocol described above.

### Immunohistochemistry Analysis

Resected tumors were prepared in formalin-fixed paraffin embedded blocks and cut in 5μm sections. Briefly, after xylene incubation, ethanol dehydration and antigen retrieval, slides were incubated for 15 minutes in 3% hydrogen peroxide to block endogenous peroxidase activity. Slides then were stained with haematoxylin and eosin (H&E), Ki76, EGFR, p-EGFR, PD-1, PD-L1, and CD3. Three drops of normal goat serum in 10ml PBS was used as blocking buffer for 1 hr at room temperature before primary antibodies incubation (Vectastatin ABC Kit; Vector Laboratories). Slides were then incubated with the following primary antibodies overnight at 4°C; Ki67 (Abcam; ab833), EGFR (Cell signaling; 4267), p-EGFR (Cell signaling; 3777), PD-1 (Cell signaling; 86163), PD-L1 (Cell signaling; 13684), and CD3 (Abcam; ab16669). The sections were developed using DAB solution (DAB substrate Kit; Vector Laboratories) and counterstained with haematoxylin. Leica DM 4000B LED microscope was used to visualize the staining and generate images at 10X magnification using LAS4.7 software. The staining was graded as low (< 30% positive cells/field), moderate (30-60% positive cells/field) or high (>60% positive cells/field). The proportion of each grade for each antigen was analyzed by Chi square test and Fisher exact test to determine significance between control and treatment groups using GraphPad Prism software. The significance was considered if *p* value < 0.05.

### Statistical Analysis

Analysis of *in vivo* experiments was performed using GraphPad Prism software, version 6.0 (San Diego California, USA). For “tumor size” and “body weight” we applied regular two-way ANOVA followed by Dunnett’s test for multiple comparisons. The survival curves were compared by Log-rank (Mantel-Cox) test. All results are expressed as mean ± S.E.M with significance level set at p ≤ 0.05.

## Acknowledgements

This work was supported by Tychon Bioscience, LLC, the National Institutes of Health R21 CA185627 and the University of Minnesota. We thank Mark Sanders and the University of Minnesota University Imaging Centers for experimental and technical support.

## Notes

### Competing Interest Statement

This study was partially funded by Tychon Biosciences, LLC, which Dr. Wagner has a financial interest in and has licensed the technology from the University of Minnesota.

