## Supplemental File for "Anti-EGFR Fibronectin Bispecific Chemically Self-Assembling Nanorings (CSANs) Induce Potent T cell Mediated Anti-Tumor Response and Downregulation of EGFR Signaling and PD-1/PD-L1 Expression"

**Bispecific Chemically Self-Assembling Nanorings (CSANs) with Engineered Fibronectins for Immunotherapy**

**Contents**

**Figure S1. Characterization and binding of E1-1DD and E4-1DD**

**Figure S2. Cross-reactivity with mouse EGFR**

**Figure S3. Internalization of E1-1DD monomer and rings**

**Figure S4. EGFR phosphorylation and MTS Assay**

**Figure S5. Trimethoprim dissociation of E1-1DD rings**

**Figure S6. H&E staining of tumor samples from the *in vivo* study**

**Figure S7. Western blot analysis and quantification of the tumor lysates**

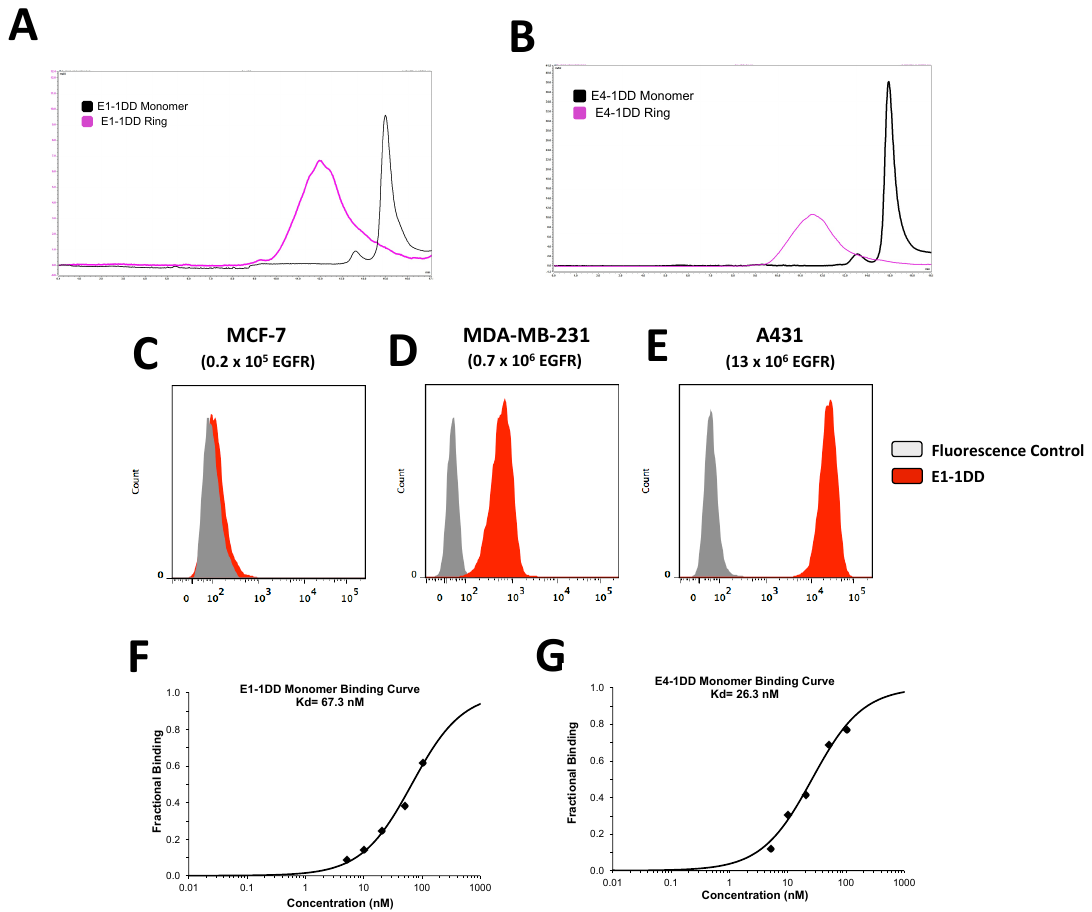

**Figure S1.**

Characterization and binding of E1-1DD and E4-1DD. **A.** SEC confirmation of ring formation and monomer purity for E1-1DD **B.** SEC confirmation of ring formation and monomer purity for E4-1DD **C.** Binding of E1-1DD rings to MCF-7 **D.** MDA-MB-231 **E.** A431 cells **F.** Affinity titration for E1-1DD monomer **G.** Affinity titration for E4-1DD monomer

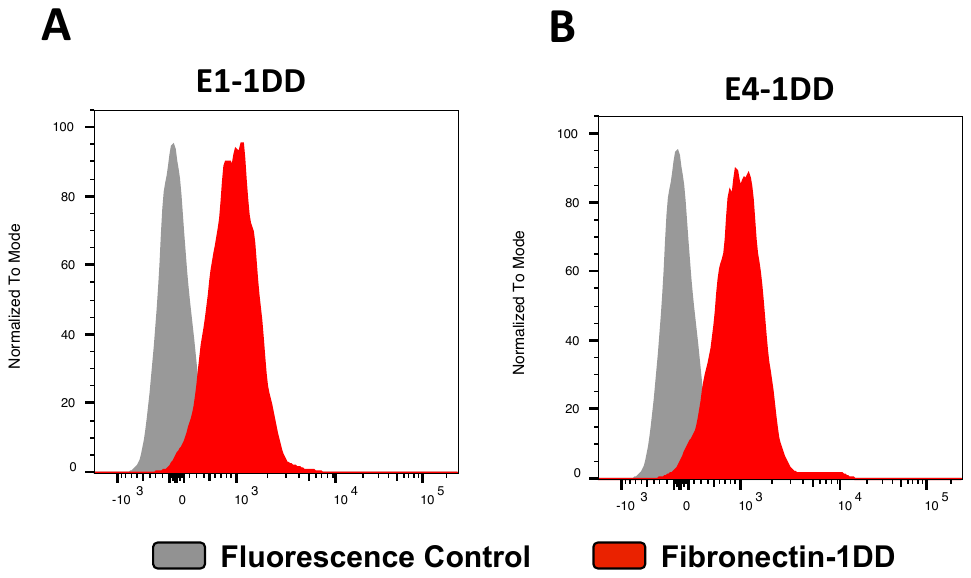

**Figure S2.**

Cross-reactivity with mouse EGFR. Binding of both **A.** E1-1DD and **B.** E4-1DD to mouse mammary cell line (HC-11) indicating these fibronectins are cross-reactive with mouse EGFR.

**Figure S3.**

Internalization of E1-1DD monomer and rings. **(A-C)** A431 cells were incubated at 4°C with **A.** PBS (Cell control) **B.** E1-1DD monomer **C.** E1-1DD ring. Both the monomer and ring show binding to the cells indicated with red. **(D-F)** After incubation of A431 cells at 4°C, samples were incubated at 37°C for 30 minutes and analyzed **D.** Cell control **E.** E1-1DD monomer internalization **F.** E1-1DD ring internalization. Red= Alexa Fluor 647-labeled E1-1DD.

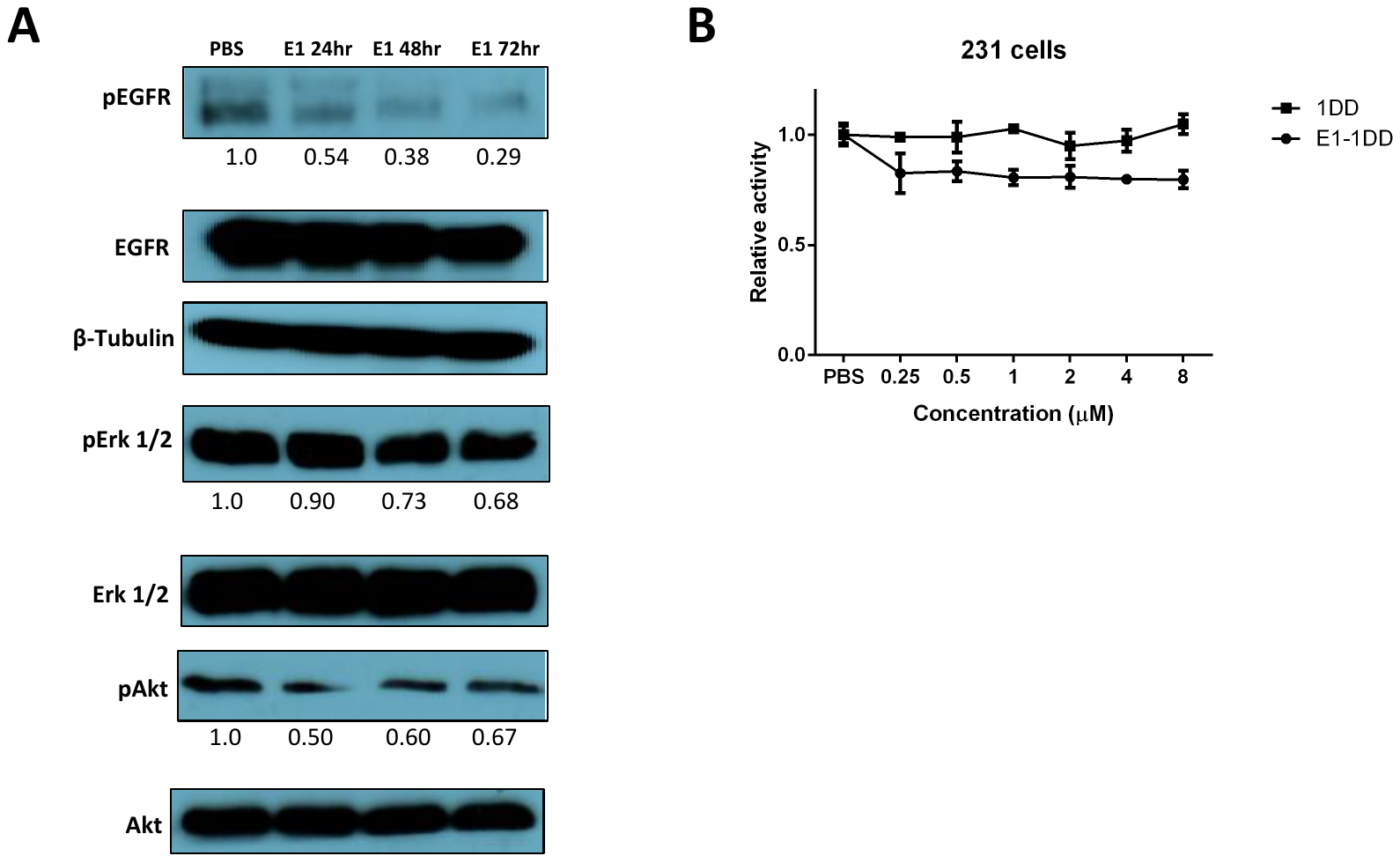

**Figure S4.**

EGFR phosphorylation and MTS Assay. **A.** Western blot analysis of E1 treated MDA-MB-231 cells. Western blot quantification by densitometry and ImageJ software. **B.** Effect of E1-1DD treatment on cell proliferation of MDA-MB-231 cells, in comparison to 1DD control, was assessed using MTS assay.

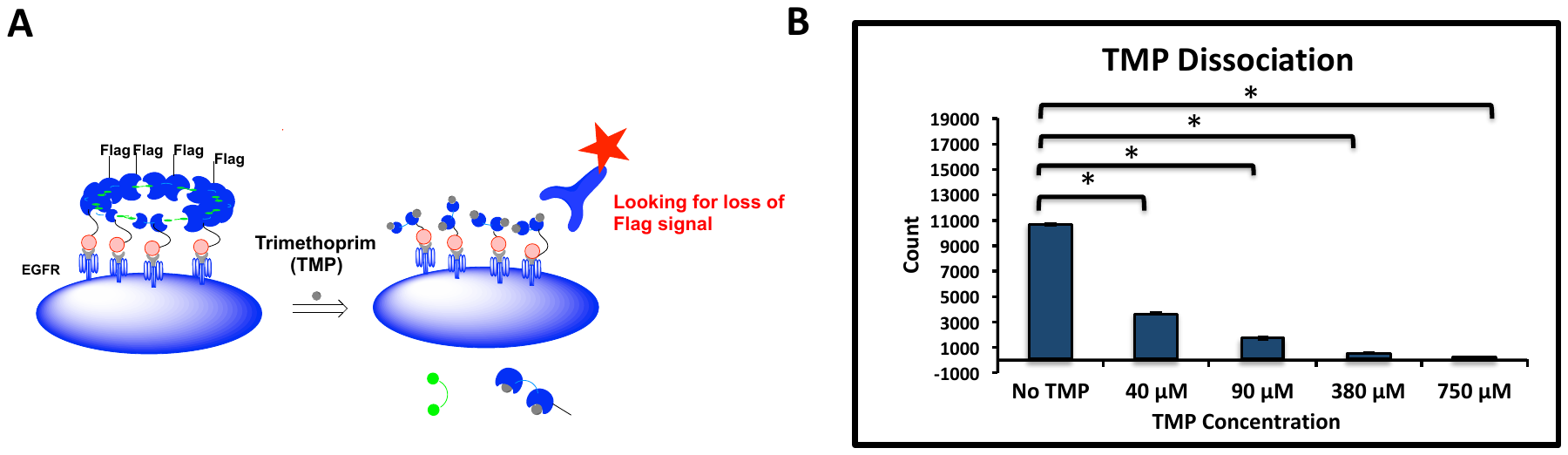

**Figure S5.**

Trimethoprim dissociation of E1-1DD rings. **A.** Schematic representation of the experiment. **B.** MDA-MB-231 cells were incubated with E1-1DD bispecific rings where 1DD has an exclusive Flag tag. Cells are incubated with the rings for an hour followed by addition of trimethoprim at different concentrations and incubated for additional 2 hours. The remaining Flag tag after this incubation is quantified with an anti-Flag Alexa Fluor 647 antibody and analyzed by flow cytometry.

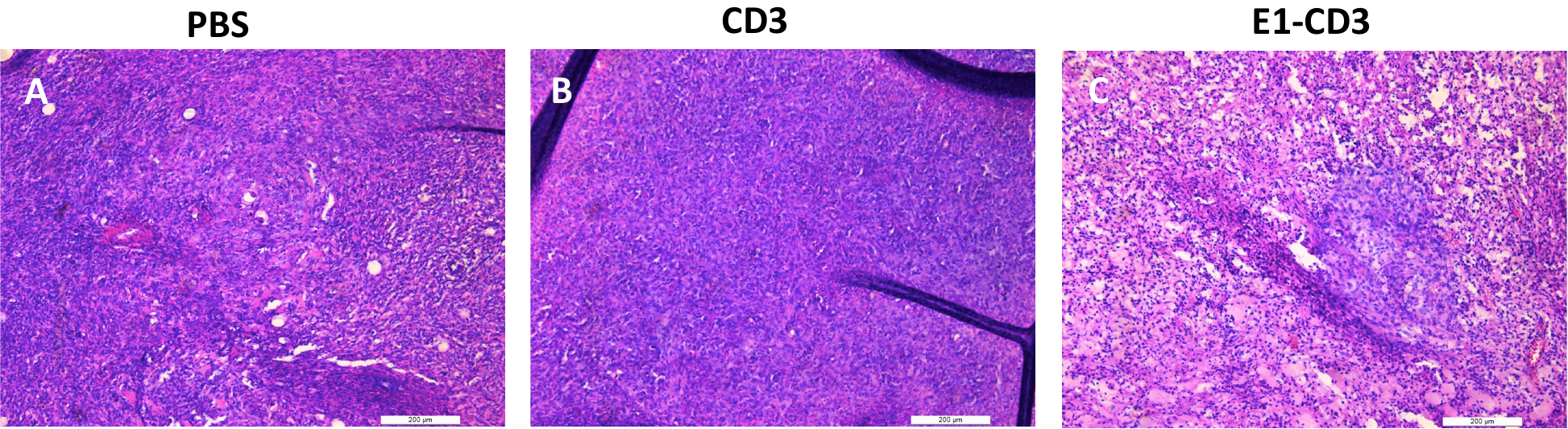

**Figure S6.**

H&E staining of tumor samples from the *in vivo* study. **A.** PBS Group **B.** CD3 Group **C.** E1-CD3 Group

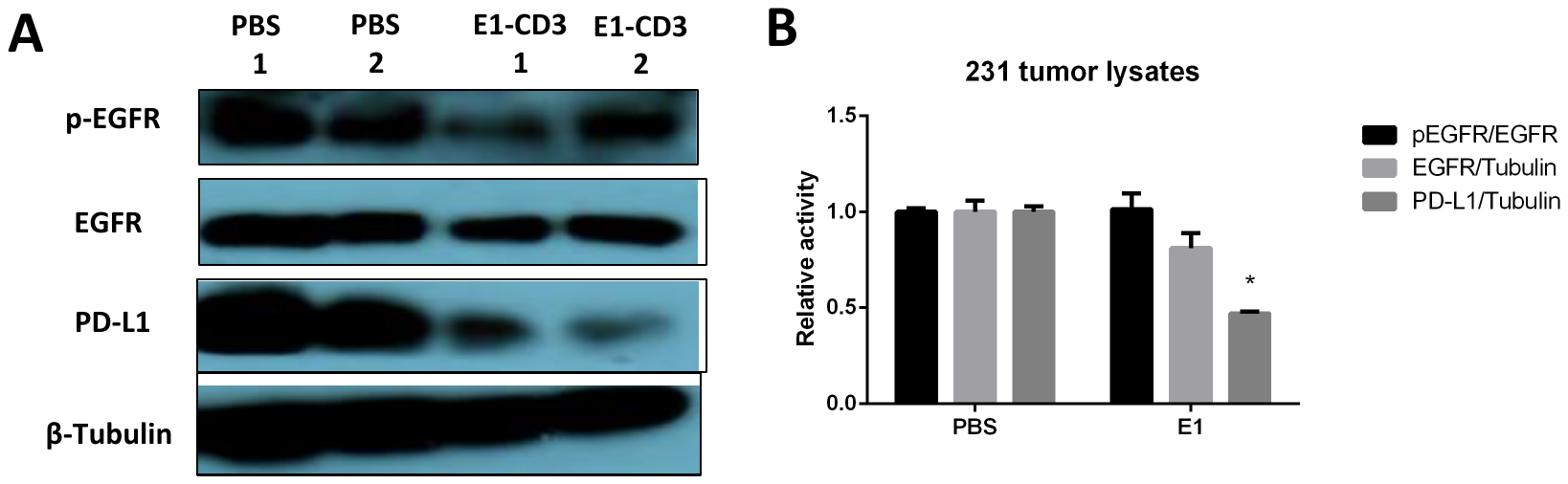

**Figure S7.**

Western blot analysis and quantification of the tumor lysates. **A.** Representative western blot analysis of tumor lysates from PBS or E1-CD3 treated mice. **B.** Western blot quantification by densitometry and ImageJ software.
